# Maturation of human intestinal epithelium from pluripotency *in vitro*

**DOI:** 10.1101/2021.09.24.460132

**Authors:** Umut Kilik, Qianhui Yu, Rene Holtackers, Makiko Seimiya, Aline Xavier da Silveira dos Santos, Barbara Treutlein, Jason R. Spence, J. Gray Camp

## Abstract

Methods to generate human intestinal tissue from pluripotent stem cells (PSCs) open new inroads into modeling intestine development and disease. However, current protocols require organoid transplantation into an immunocompromised mouse to achieve matured and differentiated epithelial cell states. Inspired by developmental reconstructions from primary tissues, we establish a regimen of inductive cues that enable stem cell maturation and epithelial differentiation entirely *in vitro*. We show that the niche factor Neuregulin1 (NRG1) promotes morphological change from proliferative epithelial cysts to matured epithelial tissue in three-dimensional cultures. Single-cell transcriptome analyses reveal differentiated epithelial cell populations, including diverse secretory and absorptive lineages. Comparison to multi-organ developmental and adult intestinal cell atlases confirm the specificity and maturation state of cell populations. Altogether, this work opens a new direction to use *in vitro* matured epithelium from human PSCs to study human intestinal epithelium development, disease, and evolution in controlled culture environments.

## Introduction

Complex three-dimensional (3D) intestinal organoids can be generated *in vitro*, modeling aspects of human tissue development, physiology, and disease (Kechele and Wells, 2019; Schutgens and Clevers, 2020). There are two major strategies for establishing organoids of the human intestine. First, intestinal stem cells (ISCs) can be isolated from primary developing or adult human gut tissues through biopsy or post-mortem sampling and propagated *in vitro* to produce epithelium-only structures, referred to as enteroids (Fujii et al., 2018; Sato et al., 2009, 2011). Culture conditions and media containing growth factor combinations have been established to facilitate ISC proliferation (Ootani et al., 2009; Sato et al., 2009) or epithelial differentiation (Fujii et al., 2018). Second, embryonic or induced pluripotent stem cells (PSCs) can be differentiated through endoderm to intestinal progenitors, which develop into immature intestinal tissues predominantly composed of epithelial and mesenchymal cells (Kechele and Wells, 2019; Spence et al., 2011; Wells and Spence, 2014). The immature organoids can then be transplanted into immunocompromised mice (Watson et al., 2014) to generate organoids with matured stem cells and differentiated epithelium with crypt-villus morphology (Finkbeiner et al., 2015; Yu et al., 2021). Combining additional germ layers or cell lineages (Eicher et al., 2021; Holloway et al., 2020; Workman et al., 2017), as well as introducing physiological triggers such as microbiota (Hill et al., 2017) or mechanical forces (Poling et al., 2018), can improve signatures of maturation. While ISC-derived intestinal organoids serve to understand homeostasis and disease conditions, PSC-derived organoids circumvent invasive tissue biopsy in humans and enable studies of development across different tissues and time points from the same reprogrammed iPSC line. However, the limited throughput and the requirement of transplantation into mice for organoid maturation limit the utility of PSC-derived organoids. In addition, it is unknown how to induce maturation of the intestinal epithelium from pluripotency *in vitro*.

Understanding the ISC niche has led to the identification of growth factors that enhance cell type composition and maturation of human small intestinal enteroid cultures (Fujii et al., 2018; Holloway et al., 2021; Jardé et al., 2020). Recent scRNA-seq studies revealed that human small intestinal epithelial development is accompanied by temporal changes of mesenchymal subtype compositions (Elmentaite et al., 2020, 2021; Fawkner-Corbett et al., 2021; Holloway et al., 2021; Yu et al., 2021). ISCs expressing Leucine-Rich Repeat Containing G Protein-Coupled Receptor 5 (LGR5) emerge concomitantly with an increase of subepithelial mesenchymal cells that express a member of the Epidermal growth factor (EGF)-family of secreted ligands, Neuregulin 1 (NRG1), during human intestine development (Yu et al., 2021). Replacing EGF with NRG1 in the tissue culture media has been shown to promote ISC maturation, organoid formation, and enteroendocrine differentiation using primary enteroid cultures and enhance ISC maturation from pluripotency (Holloway et al., 2021; Jardé et al., 2020; Yu et al., 2021). However, previous work suggested that the timing of EGF and NRG1 exposure might be important to correctly specify and mature intestinal cell fates (Yu et al., 2021). These observations prompted us to establish a novel protocol to generate differentiated human intestinal epithelium entirely *in vitro*. Here we tested the role for NRG1 by generating conventional human intestinal organoids (HIOs) from PSCs, followed by epithelial enrichment and enteroid culture in media containing Wnt3a, EGF, Noggin, R-spondin-3 (WENR), with and without NRG1. We observe that the inclusion of NRG1 in the media generates organoids from pluripotency that exhibit complex morphologies and contain enhanced diversity of intestinal secretory and absorptive epithelial cell types. Our findings suggest that further maturation of PSC-derived organoids and enteroids is possible without the requirement for transplantation into a living host and opens up possibilities to model intestinal epithelial development, disease, regeneration, and evolution from existing repositories of PSCs in a genetically tractable and high throughput manner.

## Results

Our previous findings demonstrated that PSC-derived stem cells are immature prior to transplantation into a living murine host (Yu et al., 2021). We, therefore, set out to enhance ISC maturation and differentiated epithelium from human pluripotency without transplantation (Figure 1A). We were inspired by the observation that LGR5+ stem cells and NRG1+ subepithelial mesenchyme are not observed in *in vitro* HIOs, but co-develop around the same time *in vivo* (Holloway et al., 2021; Yu et al., 2021)(Figure 1B). We first induced endoderm formation from embryonic stem cells, followed by midgut/hindgut patterning that leads to the formation of intestinal spheroids. Free-floating spheroids were embedded in Matrigel and further differentiated into HIOs in media containing EGF, Noggin, R-Spondin-1 (ENR) (Spence et al., 2011) for four weeks, leading to the co-development of epithelial and mesenchymal cells. Next, we isolated epithelium (Capeling et al., 2020; Tsai et al., 2018) and cultured epithelium-only enteroids in the presence of Wnt (WENR) for two weeks. PSC-derived enteroids were then passaged into either WENR or WENR+NRG1 media conditions to test the role of NRG1 on ISC and epithelial cell maturation. After five weeks of culture, WENR-only organoids were maintained as undifferentiated cysts (Figure 1C), whereas NRG1-grown organoids formed complex budding structures (Figure 1C). Immunostaining of NRG1-treated organoids revealed DPP4+ enterocytes, MUC2+ goblet cells, and SST+ and GCG+ enteroendocrine cells (Figure 1D). In contrast, we detected very rare or no goblet or enteroendocrine cells in WENR-only counterparts (Figure S1A).

**Figure 1.**
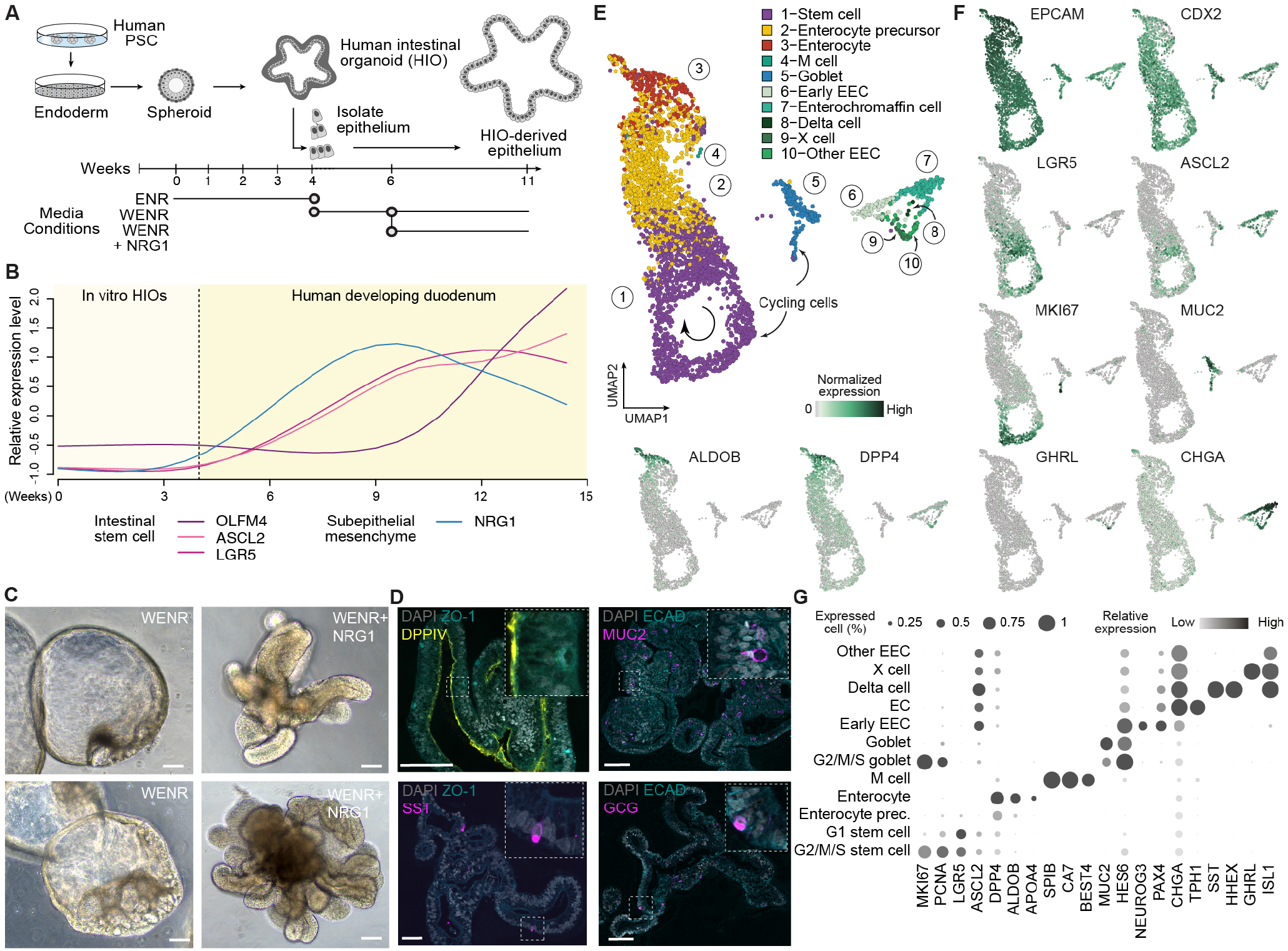
NRG1 promotes human intestinal organoid (HIO) epithelial development from pluripotency *in vitro*. (A) Schematic illustrating the generation of HIO-derived epithelium from pluripotent stem cells (PSCs). WENR stands for Wnt3a, EGF, Noggin and R-spondin-3 respectively. (B) The relative mRNA expression level of representative intestinal stem cell marker genes and subepithelial mesenchyme factor NRG1 across different time points of *in vitro* HIOs and developing duodenum based on published scRNA-seq data (Yu et al., 2021). Lines represent cubic smooth spline curves of z-transformed average gene expression levels on day 3, 7, 14, 28, and 30 *in vitro* HIOs and 8, 10, 11, 12, 14, 17, 18, 19 post-conception weeks (PCW) developing duodenum (df=5). (C) Two representative brightfield images of 11 weeks old *in vitro* HIO-derived epithelial enteroids grown in WENR (left) or WENR+NRG1 (right) conditions. Scale bar: 200um. (D) Immunofluorescence staining of 11 weeks old *in vitro* HIO-derived epithelial enteroids grown in WENR+NRG1 condition. Scale bar: 100um, DAPI: nuclei; ZO-1: tight junctions; ECAD: epithelial cell; DPP4: enterocyte; MUC2: goblet cell; SST and GCG: EEC subtypes (E) UMAP cell embedding of HIOs grown with WENR+NRG1, with cells colored and numbered by cell cluster. (F) Feature plots showing expression of selected epithelial cell type markers. (G) Dot plots showing expression of selected epithelial cell type markers.

We performed single-cell transcriptome analysis to comprehensively analyze cell heterogeneity in enteroids from each condition (Figures 1E and S1B; Table S1). Nearly all cells expressed epithelial and intestinal markers in both conditions, including EPCAM and CDX2, suggesting robust specification to the intestinal epithelial lineage. We observed striking heterogeneity in the NRG1-treated organoids and could identify at least ten molecularly distinct cell types (Figures 1E-1G; Table S2). Cell type 1 (c1) represented ISCs and highly expressed LGR5 and the ISC transcription factor ASCL2 (van der Flier et al., 2009; Schuijers et al., 2015). Interestingly, we observed the entire ISC cell cycle within c1 and the differentiation trajectory from stem cells to enterocyte precursors (c2) to mature enterocytes (c3; ALDOB+, DPP4+). In parallel, secretory lineages were present, including goblet cells (c5; MUC2+) and enteroendocrine cells (c6-c10) (Figures 1E-1G). Indeed, we observed substantial diversity in the enteroendocrine lineage, capturing precursors (c6; NEUROG3+), enterochromaffin (c7; EC; TPH1+), delta cells (c8; SST+), X cells (c9; GHRL+), and other cells (c10; ISL1+). We also detect a small population of M cells (c4; SPIB+, CA7+). On the contrary, single-cell transcriptome analysis of WENR-only organoids revealed a small proportion of enterocytes and no secretory lineages or M cells (Figure S1B; Table S2). These results indicate that NRG1 serves as an inductive, niche-derived cue to induce morphologic and cell type composition changes in enteroids derived from human PSCs.

To quantify the fidelity and maturation of cell states within WENR+NRG1 organoids, we compared organoid cells to developing and adult human reference cell atlases (Figure 2). First, we searched for nearest neighbor counterparts of the NRG1-derived cell states within a developing human endodermal cell atlas that contains hundreds of thousands of cells from the developing human lung, esophagus, liver, stomach, small intestine, and colon (Yu et al., 2021) based on single-cell transcriptome similarity (see STAR Methods) (Figure 2A and 2B). We found that WENR+NRG1 organoids consist of most similar cells to the developing human intestine (95+% map to the intestine). In contrast, WENR organoids have approximately half of the cells lacking full intestinal commitment and map to the stomach (Figures 2C and S1C). These results suggest that NRG1 may have a role in promoting or maintaining intestinal fates in the early stages of intestinal development. Next, we assessed cell type proportions of the WENR+NRG1 organoids and observed the diversity of absorptive and secretory lineages resembling the distribution of cell types from 8 post-conception-week (PCW) developing duodenum (Figure 2D). In contrast, WENR-only organoids do not possess this comparable cell-type diversity. We next assessed the maturity of WENR and WENR+NRG1 organoids by comparing single-cell transcriptomes to reference atlases of *in vitro* HIO, transplanted HIO, primary developing, and primary adult tissues (Yu et al., 2021). This analysis revealed that stem cells from WENR + NRG1 grown organoids are more mature than WENR-only cells based on their transcriptome similarity to adult ISCs (Figure 2E). Notably, WENR-only organoids are most similar to *in vitro* grown organoids and early developing duodenum ISCs (7 PCW), whereas WENR+NRG1 organoid ISCs are highly similar to transplanted HIOs (4 weeks post-transplantation) and approaching 8 PCW developing ISCs in their maturation state.

**Figure 2.**
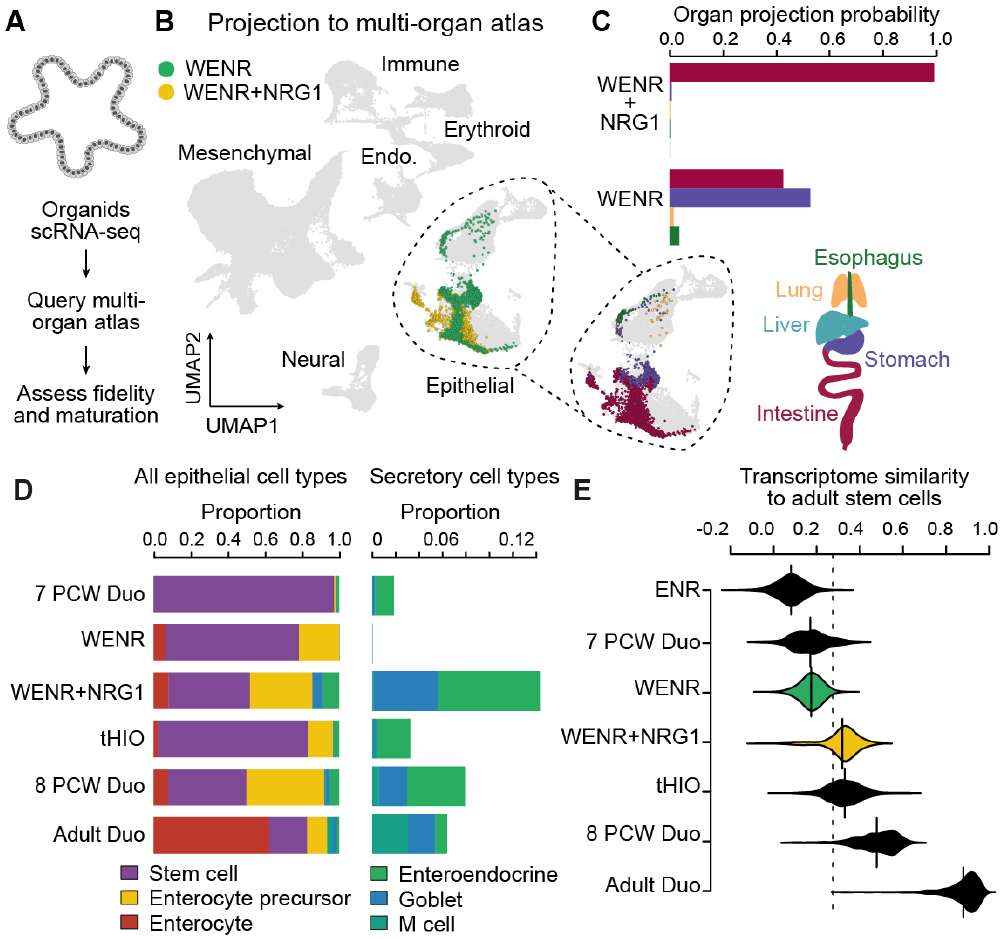
Assessment of organoid specification and maturation state through comparison to a human reference atlas. (A) HIO cells are projected onto the UMAP cell embedding of the developing human endodermal reference atlas (Yu et al., 2021). (B) HIO cells are colored by culture condition onto the UMAP of developing human endodermal cell atlas (left), zoomed into the epithelial cells to see a projection of organ identity of each cell (right). Light grey cells indicate developing human cells. The schematic represents the organ identity of developing human organs. (C) Bar plot shows the proportion of HIO cells of each culture condition mapping to each organ. (D) Stacked bar plot showing epithelial cell type composition (left) and percentage of secretory cell types (right) among epithelial cells of organoids or primary duodenum tissue. (E) Bean plot showing the distribution of transcriptome similarity (Spearman’s correlation coefficients) between adult duodenum stem cell average and stem cells of organoids, developing or adult duodenum, using the phase markers of intestinal stem cell maturation trajectory spanning from spheroids to adult stage as feature genes (Yu et al., 2021). The dashed line represents the mean of all samples. Age of the organoids: ENR, 4-week *in vitro*; WENR and WENR+NRG1, 11-week *in vitro*; tHIO, 4-week *in vitro*, and 4-week in-vivo.

Altogether, here we describe a PSC-derived enteroid protocol consisting of matured stem cells, diverse absorptive and secretory cell lineages robustly specified to the intestine, which shows symmetry breaking and primitive crypt-villus morphology. Organoid cell states are highly similar to the developing human intestine and can emerge entirely *in vitro*.

## Discussion

It has been a significant challenge to generate human intestinal epithelial organoids containing matured intestinal stem cells and differentiated epithelium from human pluripotent stem cells. Here we show that more accurately mimicking the developmental morphogens present during ISC niche development can cue stem cell maturation such that the stem cells are competent to differentiate into diverse absorptive and secretory lineages. Critically, PSC-derived enteroids grown in NRG1 bypasses the requirement for transplantation into mice for maturation, achieving a maturation of 4 week-post-transplantation organoids, allowing the PSC-derived stem cells to mature from an approximate time of developing human equivalent of 7 PCW to around 8 PCW. In addition, the enteroids are proliferative and can be passaged and expanded for high throughput experiments. Many intestinal epithelial disorders have neonatal, pediatric, or childhood onsets (Rigoli and Caruso, 2014; de Santa Barbara et al., 2002). Progress in understanding these disorders has been hindered by the inability to mature epithelium from pluripotency entirely *in vitro*. Our advancement provides exciting opportunities to use induced pluripotent stem cells for disease modeling and drug screening and may reduce the use of animal model systems in human organoid research.

Our current findings also show proof of concept that accurately mimicking a developmental niche enhances the developmental fidelity of organoid model systems and it will allow us to understand how other niche-derived factors impact naive epithelial development and maturation. Interestingly, we observed substantial enteroendocrine diversity in the presence of NRG1, and the relationship between the molecular mechanisms involved in NRG1-induced stem cell maturation and enteroendocrine diversification are unknown. In addition to NRG1, there are many other secreted molecules from mesenchyme, neurons, endothelium, and immune cells that are developmentally and spatially regulated (Holloway et al., 2021; Yu et al., 2021). It will be exciting to understand how other factors modulate the development and differentiation of each intestinal epithelial cell type in humans. Intriguingly, the stem cells and epithelium generated in this protocol are not as mature as cell states observed in adult intestinal tissues. Future experiments testing morphogen, metabolic, microbial, or other signaling pathways could build on this system to reach the ultimate goal of temporally programmed human intestinal epithelial cell states from pluripotency. More broadly, our work shows that ontogenic analysis of interlineage interactions during niche development can be leveraged to generate novel organoid culture systems.

## ACKNOWLEDGEMENTS

We thank Charlie Childs and Philipp Wahle for fruitful discussions, Dinko Pavlinic and Simone Picelli from IOB Single Cell Genomics Platform for their help with the scRNA-seq library preparation, Christian Beisel and the Genomics Facility Basel with sequencing, Diego Calabrese from University of Basel Histology Core Facility with tissue processing, Michael K. Dame from Translational Tissue Modeling Laboratory (TTML) for providing L-WRN conditioned media (Miyoshi and Stappenbeck, 2013)

## AUTHOR CONTRIBUTIONS

U.K. established the protocol and performed all experiments, with support from R.H, M.S and A.S. Q.Y. processed, analyzed, and maintained the scRNA-seq data, with initial analysis performed by U.K. U.K., Q.Y. and J.G.C prepared the figures. J.R.S and B.T provided critical reagents for this study. U.K., Q.Y, J.R.S., and J.G.C. designed the study and wrote the manuscript. All authors read and approved the manuscript.

## DATA AVAILABILITY

The accession numbers for the mRNA sequencing data and processed data reported in this paper are ArrayExpress: E-MTAB-10876.

## CODE AVAILABILITY

The code used for single-cell analysis and data presentation is being packed up,and will be deposited to GitHub soon. It is available upon request.

## STAR Methods

### Generation and culture of *in vitro* human intestinal organoids (HIO)

The use of human embryonic stem cell (ESC) line-H9 and generation of intestinal organoids were approved by the Ethics Committee of Northwest and Central Switzerland (2019-01016) and the Swiss Federal Office of Public Health. hESC line (H9) was cultured with mTESR-plus (STEMCELL Technologies) media on Matrigel (Corning, 354277) coated plates. The cells were passaged when they reached 80-90% confluency every 4-5 days. hESCs were differentiated into HIOs following the previously described protocols (Capeling et al., 2020; Spence et al., 2011; Tsai et al., 2016). In brief, hESCs were patterned into definitive endoderm by Activin A (100 ng/ml) induction in RPMI-1640 media for three days with increasing concentrations of FBS (0%, 0.2%, and 2%). Then, midgut/hindgut patterning of endoderm was induced by introducing FGF4 (500 ng/ml) and CHIR99021 (2uM) with daily media changes for up to 6 days in the presence of 2% fetal bovine serum (FBS). Spheroids were collected after 96, 120, and 144 hours of patterning and embedded in Matrigel (Corning, 354234), cultured in ENR media (mini gut basal media supplemented with EGF (RD Systems, 100 ng/ml), Noggin (Peptrotech, 100 ng/ml) and R-Spondin-1-Fc (5% conditioned media)(Ootani et al., 2009). Organoid media was changed every 3-4 days. HIOs were split once the mesenchyme outgrew Matrigel, and excessive cell debris accumulated in the core every 7-10 days. The mini gut basal media consists of DMEM/F-12, HEPES (Thermo Fisher Scientific, 11330032), and 1x B27 supplement (Thermo Scientific, 12587001). All media used in the differentiation process contain 1x Penicillin-Streptomycin (Thermo Scientific, 15140122).

### Isolation of epithelium from HIOs and culture of HIO-derived epithelium

Cold dispase achieves epithelium isolation from D28 HIOs (STEMCELL Technologies, 5U/ml), based on previously reported protocol with minor modifications (Capeling et al., 2020; Tsai et al., 2018). Briefly, one full 24-well plate of HIOs was recovered from Matrigel by pipetting and transferred into a Petri dish. Then, excess Matrigel and media were removed, and 3 ml ice-cold dispase was added on HIOs. Organoids are incubated for 30 minutes on ice, then the dispase is replaced by 3 ml 100% FBS and incubated for 15 minutes on ice. Next, 3 ml DMEM/F-12 was added on organoids and pipetted up and down for mechanical separation of epithelial and mesenchymal layers. Then, organoids were transferred to a 15 ml Falcon tube filled with DMEM/F-12 media, and three washes were performed at 400g for 5 minutes. While aspirating the media, mesenchymal content, which appears as a cloud on top of the pelleted epithelium, should be removed with the residual Matrigel. Alternatively, epithelial pieces could be picked up individually under a stereomicroscope. We choose to use the pellet to include singlet, doublet, and multiplets and increase the chance of enteroid recovery from ISCs. Next, the pellet should be triturated 50 times in the residual volume of media and then mixed with Matrigel by calculating 50 μl per single well of a 24 well plate to have 9 to 12 wells of epithelium cultured in complete enteroid media (WENR) (Tsai et al., 2018). After 10-14 days of culture, epithelial cysts were passaged and embedded in fresh Matrigel, cysts derived from the same pool of cells cultured with or without NRG1 (RD Systems, 100 ng/ml) in the WENR conditions. Next, the growth of the organoids was monitored, and they were passaged every 7-8 days. Complete enteroid media (WERN) consists of DMEM/F-12, HEPES (Thermo Fisher Scientific, 11330032), N-2 supplement (1x), B27 supplement (1x), penicillin/streptomycin (100U/ml), N-Acetyl-L-cysteine (2mM), Nicotinamide (20mM), EGF (100 ng/ml) and 50% conditioned media containing Wnt3a, Rspondin3 and Noggin at 1:1:1 ratio. For the first three days of enteroid culture, enteroid complete media should contain Thiazovivin (2.5μM), SB431542 (100nM), Y-27632 (10μM), and CHIR99021(4μM).

### Tissue processing and imaging

Organoids were dislodged from Matrigel by pipetting up and down, washed with PBS at 400g for 2 minutes. Excess PBS and Matrigel were removed as much as possible and replaced with 10% NBF. The organoids were fixed for 24 hours on the orbital shaker. Next, organoids were washed 3x with UltraPure Distilled Water (Fisher Scientific, 12060346) for 30 minutes each and embedded as a block in Histogel (Thermo Scientific, HG-4000-012) to reduce the loss during pipetting steps. Then, stepwise dehydration was accomplished by applying methanol series (25%, 50%, 75%, 100%). Before paraffin processing, tissue was equilibrated to 100% ethanol for an hour, followed by 70% ethanol. Paraffin perfusion was done by an automated tissue processor (Medite TPC 15 Duo). Next, the paraffin processed tissues were embedded and cut to sections of 5 um thickness. Paraffin sections were baked at 60 degrees for 45 minutes. Then, slides were immediately deparaffinized (HistoChoice, H2779) and rehydrated. Antigen retrieval was performed by steaming slides in a sodium citrate buffer (pH 6.0) for 20 minutes. Next, the slides were blocked with 5% normal donkey serum (Jackson ImmunoResearch, 017-000-121) in PBS with 0.1% Tween-20 for 1 hour at room temperature. Primary antibodies were diluted in blocking solution ranging between 1:100-1:200, and slides were incubated overnight at 4 degrees in a humidified chamber. The next day, slides were washed and incubated with secondary antibodies at 1:500 dilution in a blocking buffer for 1 hour at room temperature together with DAPI staining (1 μg/ml). Slides were washed with PBS and mounted using Prolong Gold (Fisher Scientific, P10144). A list of antibodies can be found in the Key Resources Table. Image acquisition was performed using Nikon Ti2 inverted microscope with CREST X-light V3 confocal spinning disc. The light source was a Celesta laser from Lumencor equipped with a Hamamatsu ORCA Flash4 V2 camera. The objective was 40x (Nikon) with Apochromat 0.95 NA.

### Organoid dissociation and single-cell RNA sequencing

HIOs were dissociated for scRNA-seq based on previously published protocol (Miller et al., 2020). Firstly, all the tubes and pipette tips were coated with 1% BSA in HBSS to prevent the adhesion of cells. Next, organoids were mechanically dislodged, pipetted up and down to remove excess Matrigel, and pooled in a Petri dish. An excess amount of media was removed, and the organoids were minced into smaller fragments in HBSS. Tissue pieces were transferred to a 5 ml conical tube containing Mix 1 from Neural Tissue Dissociation Kit (Miltenyi Biotec, 130-092-628) and incubated on a shaker for 15 minutes at room temperature. Next, Mix 2 was added to the tube, and every 10 minutes, the mix was agitated with a P1000 pipette tip. Next, cells were filtered by a 70 μm strainer and washed three times at 400g for 5min with 1% BSA in HBSS. Cells were counted with a hemocytometer and immediately continued with 10x Chromium for single-cell droplet generation. The libraries were prepared using the Next GEM Single Cell 3’ v3.1 kit according to manufacturer instructions. The libraries were sequenced at Illumina’s Novaseq 6000 system.

### Single-cell RNA sequencing (scRNA-seq) data preprocessing

We used Cell Ranger (10x Genomics) to demultiplex base call files to FASTQ files and align reads. Default alignment parameters were used to align reads to the human reference genome GRCh38 provided by Cell Ranger (version 3.0.0). Processed single-cell (sc) RNA-seq data of the reported human developing multi-organ cell atlas and adult human duodenum cell atlas were retrieved from Mendeley data: https://doi.org/10.17632/x53tts3zfr (Yu et al., 2021).

Seurat (v3.1) package (Butler et al., 2018) was applied to the scRNA-seq data for preprocessing. Cells with less than 1,000 genes or more than 100,000 Unique Molecular Identifiers (UMIs) and those with mitochondrial transcript proportions higher than 20% were excluded. Ribosomal genes, mitochondrial genes, and genes located on sex chromosomes were further removed before normalization. After log-normalization, we analyzed each dataset separately. First, 3,000 highly variable genes were identified using the default vst method. The normalized expression levels were then z-transformed, followed by principal component analysis (PCA) for dimension reduction. Finally, uniform manifold approximation and projection (UMAP)(Becht et al., 2018) was applied to the top 20 principal components (PCs) of the organoid dataset to visualize cellular heterogeneity.

### Cell type annotation

We combined de novo cell clustering and manual curation according to the reported marker gene expression pattern and query to reference atlas to perform high-resolution cell-type annotation. Firstly, we performed Louvain clustering (resolution = 1) to each dataset. We identified cluster markers using the presto package. We used the area under the receiver operator curve (AUC) > 0.6, log-transformed expression level fold change > 0.1, Benjamini-Hochberg adjusted Wilcoxon P-value < 0.05, percent of observations in the group with non-zero feature value > 20%, and the difference between the percent of observations in the group and out of the group > 20% as the significance cut-off. Based on expression patterns of canonical cell type markers reported in previous literature (LGR5, ASCL2 for stem cells; DPP4 and ALDOB for enterocytes; MUC2 for goblet; CHGA and ISL1 for enteroendocrine cells (EEC)), together with the de novo identified cluster markers as described above, we separated the de novo identified clusters into three populations, i.e., the stem-cell-to-enterocyte population, goblet cell population, and EEC population.

Next, we removed the potential doublet of secretory and non-secretory populations. We performed sub-clustering in the three sub-populations: stem cell-to-enterocyte population, goblet cells, and EEC separately. We identified cluster markers using the presto package with the significance cut-off as described above. We defined the potential doublets as the cluster showing enriched expression levels of markers from other sub-populations, higher detected gene numbers than other clusters, and random distribution in the UMAP cell embedding. Additionally, we defined a cluster with lower detected gene numbers and a lack of specific positive markers as potential low-quality cells. Both the potential doublet and low-quality cells were removed from downstream cell type analysis.

Finally, we performed manual high-resolution cell-type annotation in the stem-cell-to-enterocyte population, goblet cell population, and EEC population separately. For the stem-cell-to-enterocyte population, considering differentiation from stem cells to enterocytes is a continuum, to ensure annotation of stem cells, enterocyte precursors and enterocytes are comparable across datasets, we projected those cell types annotated in the developing duodenum reference (Yu et al., 2021) to the corresponding population of organoids. To do this, we used the highly variable genes of the stem cell-to-enterocyte population identified in the developing reference as feature genes to calculate Spearman’s correlation coefficients between each developing cell and each organoid cell. Each organoid cell was annotated as the cell type with the highest frequency among its 20 nearest developing cells. Among the annotated stem cells in the organoids, we defined the clusters with enriched expression levels of MKI67 and CDK1 as S/G2/M-phase stem cells, others as G1-phase stem cells

For the goblet cell population, we performed sub-clustering of the goblet cell cluster to annotate goblet cell sub-states and defined the cluster with an enriched expression of MKI67 and CDK1 as S/G2/M-phase goblet cells, others as G1-phase goblet cells.

For the EEC population, we found that sub-clustering could not perfectly fit with canonical subtype marker expression for annotation of EEC subtypes (Beumer et al., 2020; Gehart et al., 2019), so we performed subtype annotation based on enrichment of aggregated expression of subtype feature sets. Based on the inspection of individual reported EEC subtype marker gene expression, we found that the EEC population could be separated into three sub-groups, i.e., immature EECs (DLL1+ and NEUROG3+), enterochromaffin (EC) cells (REG4+, TAC1+, and TPH1+) and non-EC cells (ISL1+ and CHGA-low). We first identified feature sets of each subtype as the top 20 genes showing the highest Pearson’s correlation coefficient to the aggregated expression of positive markers of each sub-group as mentioned above. Then we calculated the aggregated expression levels of each feature set and used the log-transformed aggregated expression levels across cells as a subtype enrichment score. EECs are annotated as the subtypes with the highest enrichment score. Among the non-EC cells, we found a subset showing enriched expression levels of X cell marker (GHRL), or Delta cell (SST and HHEX), so we further extracted these subtypes in the non-EC population based on enriched expression levels of respective markers as mentioned above.

After high resolution cell type annotation, we identified markers of these obtained cell types using the presto package. We used the area under the receiver operator curve (AUC) > 0.6, log-transformed expression level fold change > 0.1, Benjamini-Hochberg adjusted Wilcoxon P-value < 0.05, percent of observations in the group with non-zero feature value >20%, and the difference between the percent of observations in the group and out of the group > 20% as the significance cut-off.

### Inference of organ identity of organoids

We inferred the organ identity of HIOs based on projection to human developing multi-organ reference atlas as described before (Yu et al., 2021). Firstly, we calculated Spearman’s correlation coefficient between the organoid cells and every cluster of each reference sample using the highly variable genes defined in the combined reference atlas. We then performed z-transformation to correlations of each organoid cell with different clusters of each developing sample and concatenated them as the final Reference Similarity Spectrum to the developing human atlas representation (RSS in short). To predict the organ identities of organoid cells, we identified the k-nearest (k = 20) developing human cell neighbors of each organoid cell by calculating Euclidean distance between RSS of organoid cells and the reported Cluster Similarity Spectrum (CSS) matrices of the reference (Yu et al., 2021) using the ‘nn2 function in the RANN package. The most frequent human cell identity among the nearest neighbors was defined as the mapped human *in vivo* identity of each organoid cell.

To allow the projection of the organoid cells to the UMAP embedding of the developing human multi-organ atlas, we retrieved the reported CSS-based UMAP model of the developing human atlas, which was trained by the ‘umap’ function (with ‘ret_model’ parameter as TRUE) implemented in the uwot package (Yu et al., 2021). Then, based on the RSS of organoid cells and the pre-trained UMAP model, we used the ‘umap_transform’ function in the uwot package to project organoid cells to the UMAP embedding of the reference. Both CSS matrices and UMAP model of the developing human reference atlas were retrieved from Mendeley data: https://doi.org/10.17632/x53tts3zfr (Yu et al., 2021).

## Supplemental Figures

**Figure S1.**
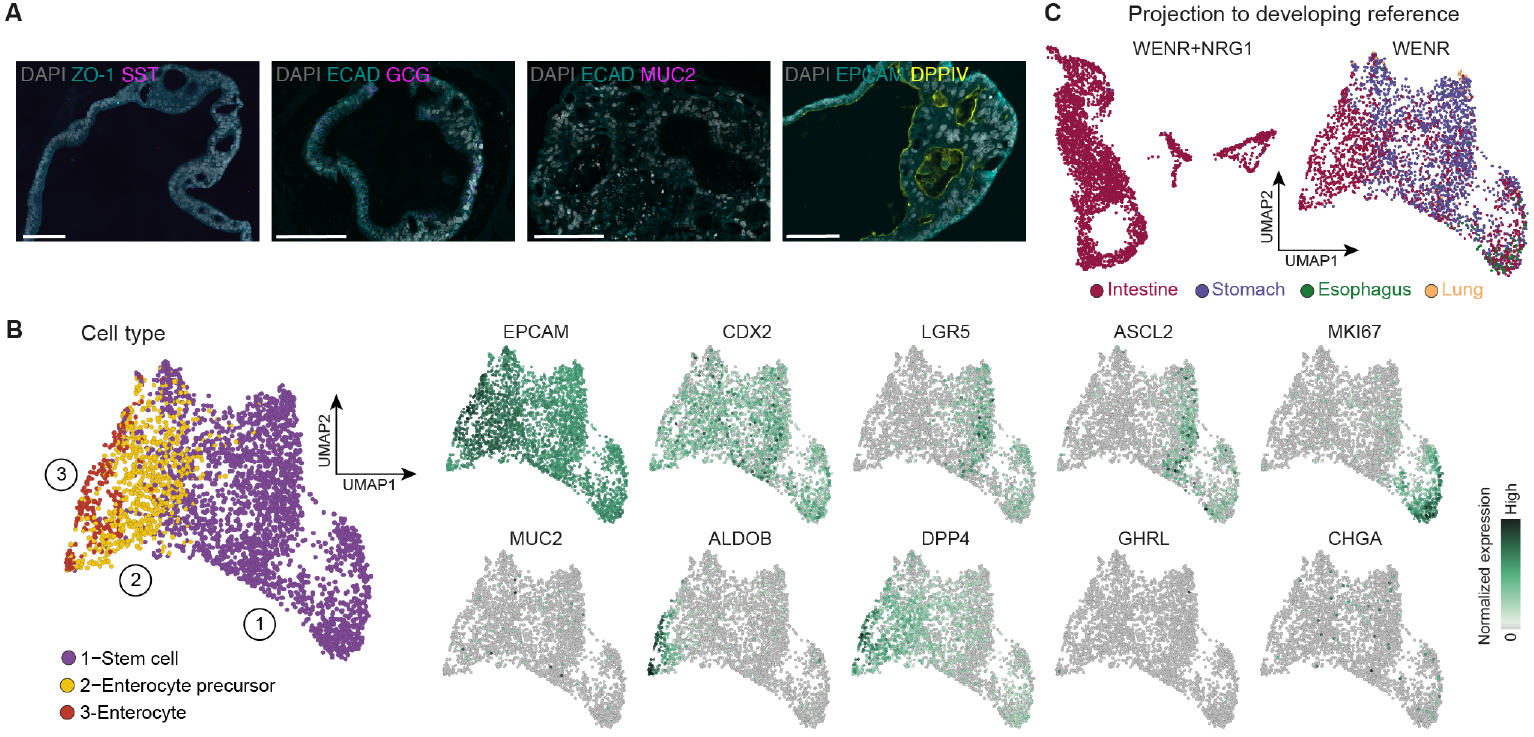
Organoid epithelium in the absence of NRG1 does not mature *in vitro*, related to Figure 1. (A) Immunohistochemistry of 4 week-HIO-derived epithelium grown for another 7 weeks in WENR conditions. Scale bar: 100um. (B) UMAP cell embedding of HIOs grown with WENR, with cells colored by cell types (left). Feature plots showing expression of selected epithelial cell type markers (right).(C) UMAP cell embedding of organoids grown with WENR+NRG1 (left) or WENR (right) media, with cells colored by projected developing human organ identity.

## Supplemental Tables

**Table S1: Sample information and scRNA-seq data quality check, related to Figures 1 and S1..**

This table presents sample information and cell number, median detected gene number, medium UMI number, and medium percentage of transcripts mapped to the mitochondrial genes of each sample after filtering.

**Table S2: Cell type markers of organoids grown in WENR or WENR+NRG1 media, related to Figures 1 and S1..**

The table presents the top 100 cell type markers (ranked by AUC) of organoids sequenced for this study.

